# CriSNPr: a single interface for the curated and de-novo design of gRNAs for CRISPR diagnostics using diverse Cas systems

**DOI:** 10.1101/2022.02.17.479653

**Authors:** Asgar Hussain Ansari, Manoj Kumar, Sajal Sarkar, Souvik Maiti, Debojyoti Chakraborty

## Abstract

Nucleic acid detection and variant calling through CRISPR-based diagnostics (CRISPRDx) has facilitated clinical decision-making, particularly during the COVID-19 pandemic. This has been further accelerated through the discovery of newer and engineered CRISPR effectors, expanding the portfolio of such diagnostic applications to a wide variety of pathogenic and non-pathogenic conditions. However, each diagnostic CRISPR pipeline requires customized detection schemes originating from fundamental principles of the Cas protein used, its guide RNA (gRNA) design parameters, and the assay readout. This is particularly relevant for variant detection, an attractive low-cost alternative to sequencing-based approaches for which no *in silico* pipeline for the ready-to-use design of CRISPR-based diagnostics currently exists. In this manuscript, we fill this lacuna using a unified webserver CriSNPr (CRISPR based SNP recognition), which provides the user the opportunity to de-novo design gRNAs based on six CRISPRDx proteins of choice (*Fn/enFn*Cas9, *Lw*Cas13a, *Lb*Cas12a, *Aa*Cas12b, and Cas14a) and query for ready-to-use oligonucleotide sequences for validation on relevant samples. In addition, we provide a database of curated pre-designed gRNAs and target/off-target for all human and SARS-CoV-2 variants reported so far. CriSNPr has been validated on multiple Cas proteins and highlights its broad and immediate scope of utilization across multiple detection platforms. CriSNPr is available at URL http://crisnpr.igib.res.in/.

## Introduction

Highly specific recognition of DNA and RNA by CRISPR proteins has made them useful not only as primary gene editors but also for rapid molecular diagnosis of pathogenic mutations in nucleic acids. Traditional probe-based diagnostic tests rely on a polynucleotide hybridizing to the target DNA/RNA and providing a readout through an amplification reaction (1–4). Such quantitative real-time polymerase chain reaction (qRT-PCR) tests and their derivatives have proven to be the gold standard for detecting trace amounts of pathogenic nucleic acids in samples and have been used worldwide for detecting SARS-CoV-2 during the ongoing COVID-19 pandemic. Although qRT-PCR is highly sensitive to detecting only a few copies of the target pathogenic sequence, its ability to distinguish very closely related sequences has not been successfully demonstrated for robust clinical diagnosis to date.

CRISPR-based diagnostic approaches rely on the association of Cas proteins with a target nucleic acid followed by a secondary readout either directly through the bound ternary complex or catalytic cleavage of the substrate and subsequent collateral cleavage of reporter molecules (5–10). The steps of DNA/RNA interrogation involve guide RNA (gRNA) binding followed by the catalytic activity of Cas effectors. It has been demonstrated that for some of the Cas proteins, this two-step process generates a very high specificity of target recognition that can be extended for the diagnosis of single nucleotide variants (SNVs) (11–20). Contrasted with the gold standard SNV detection technologies relying on Sanger/Deep sequencing which require dedicated infrastructure, manpower, and analysis pipelines coupled with longer turnaround times, CRISPR-based variant calling is an attractive alternative for rapid, low-cost diagnosis of disease-causing mutations. Currently (till January 2022) the ClinVar database lists about 117437 (GRCh38) pathogenic human variants associated with diseases, a large majority of which can be detected through CRISPR-based tests (16–18,20–23). Similarly, the rapidly evolving SARS-CoV-2 variants highlight the importance of detecting mutations in pathogenic sequences for developing public health strategies, effective vaccines, and understanding disease pathophysiology.

CRISPRDx is a relatively recent addition to the repertoire of diagnostic methodologies for detecting SNVs. The majority of these pipelines rely on the ability of the Cas protein to distinguish nucleic acids based on mismatches in the gRNA at defined positions. Such nucleotide position-specific mismatch sensitivity which was first reported with *Lw*Cas13a has now been shown to be associated with several other Cas effectors and employed for SNV detection, such as *Fn/enFn*Cas9, *LbCas12a, AaCas12b*, and Cas14a (14–20,24). Several CRISPR/Cas systems have also demonstrated Protospacer Adjacent Motif (PAM) mismatch sensitivity but as PAM is not always present at the target DNA/RNA sequences, their applicability for diagnostics assays is limited (16,25).

Although the overall approach for mismatch identification remains largely similar across the Cas proteins, each individual Cas protein has unique properties with respect to mismatch-sensitive positions in the gRNA. These have been identified by detailed nucleic-acid: protein structural and biochemical studies. Thus, while the same SNV can be targeted by multiple Cas proteins, each diagnostic strategy requires an individualized crRNA design (14–20,24). Considering that every Cas system is different in terms of its gRNA sequence, readout mode, PAM requirement, and mismatch sensitivity positions, it can be time-consuming for a user to first identify which suitable Cas effector to use and then design respective gRNA and primers for performing diagnostic assays. Thus, it is necessary to develop a unified approach that can help any user with minimum knowledge and information required for designing such detection assays for any SNV of interest.

To address this, we report a web server named CRISPR-based SNP recognition (CriSNPr) for designing CRISPR-based diagnostics pipelines across the CRISPR platforms reported so far for variant detection. CriSNPr provides a pipeline for CRISPR-based detection of pathogenic and non-pathogenic mutations for all reported human nucleotide variants (dbSNP) and SARS-CoV-2 including variants of interest/concern (VOI/VOC). In addition, it allows the design and implementation of *de novo* variants of choice. The server queries for the SNV of interest and provides users with information regarding all Cas systems that can be used to detect that particular SNV, and the required crRNA and primer design parameters based on gRNA design principles available in the literature for each Cas protein. Importantly, CriSNPr, unlike other available sgRNA designing tools, also gives information about the off-targets for the SNV targeting modified crRNA sequences.

CriSNPr has integrated existing information about mismatch sensitive positions for *Fn/enFn*Cas9, *Lw*Cas13a, *Lb*Cas12a, *AaC*as12b, and Cas14a and has been experimentally validated on SNVs for a subset of the Cas proteins successfully (14–20,24). To expand its immediate application towards existing human and SARS-CoV-2 variants, the current version of CriSNPr contains SNV information from human dbSNP and SARS-CoV-2 CNCB-NGDC respective databases, such that even with no prior sequence information about an SNV, a user can avail designed sequences by using the dbSNP rsID for humans or mutant amino acid position for SARS-CoV-2 respectively (26,27). For new SNV containing sequences that are not previously available in CriSNPr’s database, users can fetch crRNA and primer sequences by providing a sequence length of 20-30 nucleotides with SNV position and variant nucleobase identity.

## METHODS

### Oligos

A list of all oligos (Merck) used in the study can be found in **Supplementary Table 1**.

### Generation of CriSNPr database

#### For the human SNPs/SNVs with pathological relevance

To build the CriSNPr database for variant detection through various Cas systems for the population-specific SNPs and pathological mutations for humans, the latest dbSNP database build 155 (GCF_000001405.39) was downloaded from the NCBI FTP site (26). Since the focus was only to detect SNPs/SNVs all the other variation types like, insertion, deletion, duplication, and translocation were filtered-out, keeping only the variants with common SNV (Single nucleotide variant) tag and valid ClinVar ID were retained for further analysis (21,22). Over these variants, vcflib (v1.0.0) was used to convert vcf into a tabular format for smooth analysis while retaining trivial information such as chromosome coordinates, reference, and alternate alleles, Reference SNP (rsID), gene, disease, and population level’s frequency, etc (28). Additionally, multi-allelic SNPs were transformed into multiple entries for better processing and were labeled as ‘Not available’, if missing, or ‘Not provided’, for unavailable information. All the previous data filtrations and curations were performed with the help of Pandas python library (v1.3.5) (29). Finally, to design the gRNAs for detection, filtered SNVs/SNPs positions were mapped to the human reference genome (Gencode GRCh38.p13) to cross-check the reference base lies within the PAM proximity and followed by incorporation of a synthetic mismatch at positions based on the use of different Cas systems, **Figure 1** (30–32). Further, genome coordinates for the target SNV were used to acquire flanking sequences to design the primers for target amplification. Since *Fn*Cas9 does not show collateral activity, primers were designed in such a way that *in-vitro* cleaved products after enzymatic treatment can be optimally resolved, keeping a difference of 1:3 and 2:3 between the length of the cleaved products.

**Figure 1.**
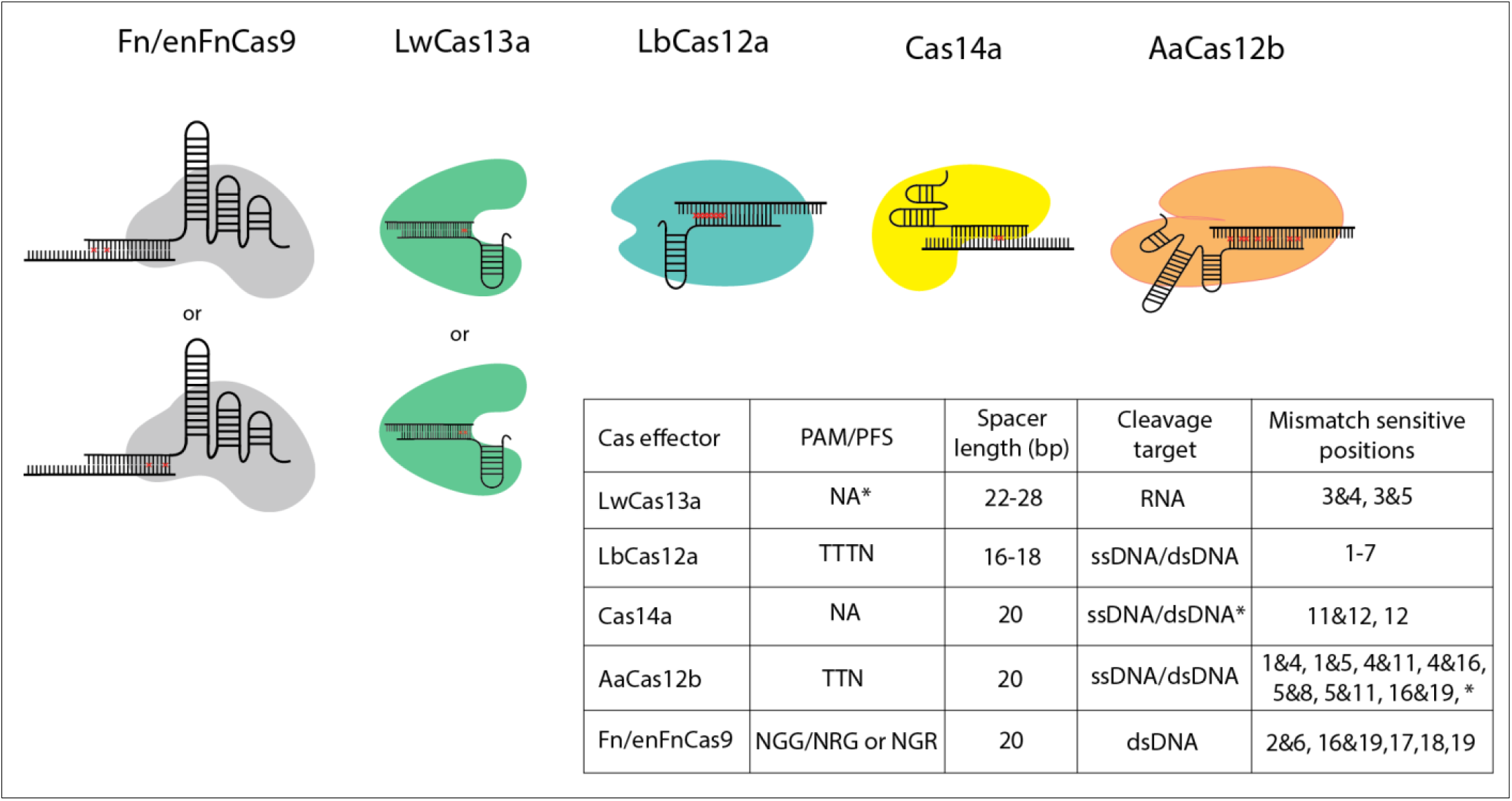
Nucleotide mismatch sensitive crRNA positions reported for different Cas systems, which are included in CriSNPr for *Lw*Cas13a, *Lb*Cas12a, Cas14a, *Aa*Cas12b and *Fn*/en*Fn*Cas9. Table summarizing the Protospacer adjacent motif (PAM)/Protospacer Flanking Site (PFS) is shown for every Cas protein along with mismatched sensitive positions reported in the literature. \**Lw*Cas13a doesn’t require PFS when targeting the mammalian genome. *Cas14a doesn’t need any PAM for ssDNA cleavage but cleavage of dsDNA is constrained by TTTA PAM. \**Aa*Cas12b have shown with mismatch sensitivity for some other nucleotide positions as well but not included here because the discrimination between WT/Mutant is not well enough.

This was achieved through the BioPython SeqIO library (v1.76) for genome parsing followed by PAM allocation using the regex library (v2021.11.10) (33). Nucleotide base incorporation was done by creating a customized function. The complete CriSNPr database was generated in the SQLite3 library (v3.31.1) of Python (v3.7.6) (34,35). The population-level SNP’s frequency data gets plotted as a bar plot using matplotlib library (v3.1.3) in real-time (36). A custom python function utilizing the Primer3 library (v0.6.1) is written to design amplicon primers (37).

#### For the SARS-CoV-2 variants

Nucleotide level variant annotation data was downloaded from CNCB-NGDC (China National Center for Bioinformation-The National Genomics Data Center), providing the latest variant information based on the analysis of the GISAID genome sequences dataset (27,38). After downloading the variant annotation data table, gene and transcript mutations were taken forward for the analysis, and variations lying in the intergenic and untranslated regions were removed. Subsequently, non-SNV mutations were also filtered from the data. Useful information such as chromosome, position, reference allele, alternate allele, amino acid change, and variation frequency was included for further analysis. The number of virus sequences with an SNP is plotted as a bar plot using matplotlib library in real-time (36). The standard CoV-2 variation nomenclature requires a gene name followed by a “_” (underscore), reference amino acid, position at protein, and alternative amino acid. Accordingly, a column was added for the variation identity which could be used as a query by the user. gRNA and primers were designed as described earlier for the human dbSNPs dataset.

### Generation of Seq-CriSNPr

CriSNPr provides sequence-based variant detection in real-time by designing crRNA and primers. For this, the user needs to enter sequence along with variant position and identity. At first, variant base identity was distinguished with respect to the reference base at the user-provided position. If variant base identity is the same as the reference base, the back-end server returns an error message. An input sequence is also validated for the invalid base (non-‘ACGT’ characters) and length (20-30 nucleotides) through Flask-WTForms (v2.2.1). Next, the sequence is mapped to the user-selected organism reference genome using BWA aligner with no mismatch to fetch the variant position on the genome (39,40). If the sequence is not found in the genome as defined by the user, it is returned with a warning. Further, the genome position of the variant is queried to the existing CriSNPr database, if not found, PAM sequences are searched in near proximity as defined, **Figure 1a**. Upon PAM location, the variant base is incorporated into the sequence followed by a random mismatch at the appropriate position, **Figure 1a**. BEDTools (v2.29.2) is used to extract the gene annotation of the crRNA location (41). Next, primers were designed using the Primer3 python library (v0.6.1) as defined earlier (37).

### Off-target prediction of modified gRNAs

CriSNPr creates an off-targets link for each of the crRNA designs concurrently. Upon clicking the link, based on organism and Cas-system, a request is sent to the back-end and a file is generated in a format as required by Cas-OFFinder (v2.4) (42). Next, a customized function is created to generate file format compatible with the Cas-OFFinder and predict off-targets up to 4 mismatches by Cas-OFFinder stand-alone version in CPU mode. These off-targets are sorted ascending based on the number of mismatches to the crRNA sequence.

### Development of CriSNPr web server

Web-server was designed and developed in Flask (v1.1.1) coupled with Jquery (v3.5.1) and Bootstrap (v5.0.2) (43). All codes were written in the Python programming language (v3.7.6) and maintained through the conda environment (v4.11.0) (34,44). The full source code of the pipeline is available at https://github.com/asgarhussain/CriSNPr.

### Percentage targetability of SNVs in human and SARS-CoV-2 by individual Cas systems

CriSNPr database has crRNA for various that can be used for SNP detection. Targetability of various Cas systems was calculated with the help of CriSNPr database in Pandas library and plotted in R (v4.0.5) using ggpubr (v0.4.0) (45). Different mutation classes of various Cas systems were also analyzed in Pandas (v1.3.5) and plotted with the ggpubr R package, **Figure 5 (Supplementary Table 3 and 4)** (29,45). The intersections among the target variations from different Cas systems have been calculated using Upset modules from Intervene offline version (v0.6.5) (46,47).

### Protein purification

A plasmid containing dFnCas9 (dead or catalytically-inactive) (19) was transformed and expressed in Escherichia coli Rosetta 2 (DE3) (Novagen). Transformed Rosetta 2 (DE3) cells were cultured at 37°C in LB medium containing 50 mg/ml Kanamycin until OD600 reached 0.6. After the protein expression induced using 0.5 mM isopropyl b-D-thiogalactopyranoside (IPTG), the culture was grown overnight at 18°C. Cultured E.coli cells were harvested by centrifugation and lysed through sonication in a buffer (20 mM HEPES, pH 7.5, 500 mM NaCl, 5% glycerol and 100 mg/ml lysozyme) supplemented with 1X PIC (Roche). Subsequently, the supernatant obtained after centrifugation was put through Ni-NTA beads (Roche), and elution was performed with a buffer (20 mM HEPES, pH 7.5, 300 mM Imidazole, 500mM NaCl). The obtained elute fractions were concentrated and the protein was further purified by size-exclusion chromatography on HiLoad Super-dex 200 16/60 column (GE Healthcare). Finally, the protein was quantified by the Pierce BCA protein assay kit (Thermo Fisher Scientific) and stored in a buffer solution (20 mM HEPES pH 7.5, 150 mM KCl, 10% glycerol, 1 mM DTT) at −80°C until its use in a reaction, **Supplementary Figure 1**.

Proteins AaCas12b and Cas14a were purified as described in previously published protocols with any needed optimizations (16,18). Plasmids containing *Aa*Cas12b (Addgene no. 113433) and Cas14a (Addgene no. 112500) were transformed and expressed in Escherichia coli BL21 (DE3) (Novagen) respectively. Transformed BL21 (DE3) cells were cultured at 37°C in terrific broth (TB) medium with respective antibiotic selections and induced with 0.5 mM isopropyl b-D-thiogalactopyranoside (IPTG) upon OD600 reached 0.6. Overnight grown cultures at 18°C were harvested and cells were lysed with sonication in a buffer (50mM Tris-HCl, pH 7.5, 5mM Imidazole, and 500mM NaCl) supplemented with 1X PIC (Roche). After centrifugation, the supernatant was put through Ni-NTA beads (Roche) and washed with a wash buffer (50mM Tris-HCl, pH 7.5, 20 mM Imidazole, and 500mM NaCl). The proteins were then eluted with a buffer (50mM Tris-HCl, pH 7.5, 300 mM Imidazole, 500mM NaCl), and MBP and His tag were removed by overnight TEV protease incubation at 4°C. The filtered proteins were further purified by size-exclusion chromatography on HiLoad Super-dex 200 16/60 column (GE Healthcare) and analyzed using SDS-PAGE. After quantification by the Pierce BCA protein assay kit (Thermo Fisher Scientific) the purified *Aa*Cas12b and Cas14a were stored in a buffer (20mM Tris-HCl, pH 7.5, 250mM NaCl, 5%Glycerol, 1mM DTT) at −80°C until further use, **Supplementary Figure 1**.

### *SARS-CoV-2 variation detection using CriSNPr designed gRNAs for Aa*Cas12b, Cas14a *and Fn*Cas9

#### *Fn*Cas9 – RAY

A region from the SARS-CoV-2 S gene containing E484K mutation was used for modified gRNA and primers to be designed by CriSNPr. Reverse-transcribed and 5’ end biotin-labeled amplicons with/without mutation were used as target sequences. The chimeric gRNA was prepared by equimolar mixing of crRNA and synthetic 3′-FAM-labeled TracrRNA in a buffer (100 mM NaCl, 50 mM Tris-HCl pH 8 and 1 mM MgCl_2_) and heating at 95ºC for 2–5 min followed by slow-cooling for 15–20 min at RT. Subsequently, equimolar gRNA:dFnCas9 RNP complexes was prepared in a buffer (20 mM HEPES, pH7.5, 150 mM KCl, 1 mM DTT, 10% glycerol, 10 mM MgCl_2_) and incubated for 10 min at RT. Finally, the target 5’ end biotin-labeled amplicons were treated with the RNP complexes at 37ºC for 10 min and the readout was obtained by adding 80 μl of dipstick buffer along with a Milenia HybriDetect 1 lateral flow assay strip for 2-5 min at room temperature followed by visual or smartphone app-based (True Outcome Predicted via Strip Evaluation, TOPSE) quantification (17,20).

#### *Aa*Cas12b – Cdetection

*Aa*Cas12b based FQ detection was performed with active RNP, prepared by equimolar mixing and incubation of *Aa*Cas12b and gRNA in a buffer (40 mM Tris-HCl, pH 7.5, 60 mM NaCl, 6 mM MgCl_2_) for 10 min at RT. Subsequently, ssDNA target (60nt WT/E484K SARS-CoV-2) sequences mixed having background human genomic DNA along with custom synthesized homopolymer (poly T) 5nt ssDNA FQ reporter molecules (GenScript) were added to the reaction within a well Corning 96-well flat-bottom black clear bottom microplate. Reactions were incubated at 37ºC for the indicated time periods (up to 180min) in a fluorescence plate reader (Tecan). With fluorescence intensity measured every 10 min (λ_ex_: 490nm; λ_em_: 520nm, transmission gain: optimal), the resulting data after background subtraction using intensity values recorded in the absence of ssDNA target sequences were plotted by using an R script (16).

#### Cas14a – DETECTR

The active Cas14a RNP complexes used for FQ detection were prepared by equimolar mixing of Cas14a and sgRNA within a buffer (40 mM Tris-HCl, pH 7.5, 60 mM NaCl, 6 mM MgCl_2_) and incubating for 10 min at RT. Further, the fluorescence readouts were developed with the addition ssDNA (60nt WT/E484K SARS-CoV-2) target sequences having background human genomic DNA along with 200nM 12nt poly T ssDNA FQ reporter molecules (GenScript) into the reaction mix in a Corning 96-well flat-bottom black clear bottom microplate. Reactions were performed at 37ºC for indicated time periods (up to 180min) in a fluorescence plate reader instrument (Tecan). Fluorescence intensity was recorded every 10 min (λ_ex_: 490 nm; λ_em_: 520nm, transmission gain: optimal). Finally, the resulting data calculated after background subtraction (intensity in the absence of ssDNA target sequences) was plotted using an R script (18).

## Results

### Conventional gRNA design tools are not tailored for CRISPR diagnostic pipelines

While there are multiple *in silico* pipelines for gRNA design suited for individual Cas proteins, these are primarily intended for gene editing applications (48–57). For a given Cas9, the workflow for gRNA design encompasses integrating specificity and sensitivity scores generated from multiple factors such as identification of available PAM sites relevant to the target, presence of preferred nucleotides in proximity to PAM, their context within the full-length gRNA sequence, GC content of the gRNA and the overall correlation between gRNA sequence and experimentally validated editing rates (48–50,52,54). In addition to gRNA sequence features, the local genetic and epigenetic characteristics, presence or absence of structural motifs, nucleosome positioning, etc. are some of the factors considered for optimal gRNA design (49–51,55). Assigning specificity scores to gRNAs is done either through alignment to the genome or through assigning scores based on hypothesis-driven approaches that integrate gRNA structural information in addition to sequence data. In recent times, machine learning has also contributed to optimal computational design parameters that empirical sequence and structure-driven gRNA prediction algorithms are unable to integrate (49,55,57).

Importantly, these algorithms are designed to perform predictions that are meant to identify the most optimal gene targeting sgRNAs relying on the biological parameters of DNA recognition by the individual Cas systems. In the case of CRISPR diagnostics, the paradigm is empirically shifted-the target sgRNA binding region is relatively fixed, and diagnostic sgRNAs and corresponding amplicon or reporters need to be tailored around the target region. For every Cas used in the diagnosis of a given SNV, the accessible PAMs and the guidelines for mismatch sensitivity positions are unique. Also different are the off-targeting propensity of the gRNAs designed based on the chosen Cas effector, **Figure 1**.

For example, mismatch sensitivity associated with RNA targeting *Lw*Cas13a works in combination by placing an SNV along with a synthetic mutation either at 3rd and 4th or 3rd and 5th PAM proximal nucleotide positions within a crRNA sequence (15). In accordance with recent reports, 5’ Protospacer flanking sites (PFS) are not essential for *Lw*Cas13a when targeting mammalian sequences but ‘H’ (A/C/T) at 5’ sites could improve target binding or cleavage (15,58). Unlike *Lw*Cas13a, *Lb*Cas12a requires TTTN PAM sequence and shows sensitivity at multiple single nucleotide positions from starting from 1st base to 7th base proximal to PAM (11,14,59). As previously reported, the mismatch shows sensitivity when placed anywhere in the region of the 1st-7th base positions using ssDNA or dsDNA as cleavage targets. This difference in signals between wild-type and mutated sequences can further be improved when shorter crRNAs of length 16-18 nt were used (14). Similar to *Lb*Cas12a, Cas14a can show cleavage on both ssDNA as well as dsDNA targets sequence but when targeting dsDNA Cas14a does require a T-rich 5’ TTTA PAM while there is no such PAM constraint with ssDNA. Therefore, even though Cas14a shows sensitivity on just 11&12 or 12 crRNA positions, this PAM flexibility is a proposed advantage when used for SNV detection (13,18). Although DNA cleaving *Aa*Cas12b requires 5’ TTN PAM to cleave the ssDNA/dsDNA sequences it shows dual mismatch sensitivity at several positions increasing the possible target SNV combinations (12,16). As this dual mismatch sensitivity by *Aa*Cas12b is shown at various crRNA positions, only combinations shown with efficient wild type and mutated sequences discrimination include 1&4, 1&5, 4&11, 4&16, 5&8, 5&11 or 16&19 were selected (16). Similarly, some of our previous reports suggested *Fn*/en*Fn*Cas9 systems with dual mismatch sensitivity require mismatches at particular positions such as 2&6 or 16&19, abrogating target dsDNA binding for variant detection (17,19,20). Along with dual mismatch sensitivity, *Fn*/en*Fn*Cas9 are also recently reported to show single mismatch sensitivity at PAM distal 17th,18th, and 19th positions (24). Having the possibility to faithfully discriminate between sequences contingent single nucleotide mutations through mismatch sensitivity across the crRNA sequence provides an opportunity to repurpose and utilize all these Cas effectors for rapid diagnosis of SNVs associated with pathological or non-pathological conditions.

It is worth noting that we have included *Lw*Cas13a in our web server as it has been previously reported for SNV detection. *Lw*Cas13a belongs to a class of RNA targeting Cas effectors which do not have any PAM constraint. Thus their targetability across the genome cannot be directly compared with other DNA targeting Cas effectors, which are PAM dependent. However, designing diagnostic assays with *Lw*Cas13a includes an additional step of converting DNA to RNA through *in vitro* transcription (15).

### CriSNPR generates readouts querying variants in publicly available human and SARS-CoV-2 datasets

We selected the Single Nucleotide Polymorphism database (dbSNP) of NCBI as a *bona fide* repository of SNPs with human clinical relevance from one of the largest variation databases of humans (26). This comprises single nucleotide variations, insertions, deletions, and microsatellites juxtaposed with population-level frequency, publication, genomic annotation of common variations, and pathological mutations. We first filtered common pathological SNPs from the latest dbSNP Build 155 (**Figure 2, Methods)**, by removing uncommon, non-clinical, and non-SNP mutations. For SARS-CoV2, we filtered SNVs by removing UTR or non-SNP mutations from the well updated CNCB-NGDC SARS-CoV-2 variations database (27). For target SNVs/SNPs, crRNAs were designed by keeping variant bases at designated positions followed by synthetic mismatch at positions relevant to experimentally validated data obtained from the diverse CRISPR systems, **Figure 2**. Next, the genome coordinates of these crRNA sequences were used to obtain flanking sequences and subsequently design primer sequences through the Primer3plus python library for PCR amplicons that can be used either binding or cleavage based reporter output assays, depending on the CRISPR proteins of choice (37). To reduce the non-specific amplification by primer sequences, these were further filtered for off-targets on a representative bacterial genome database (NCBI), virus genome database (NCBI), and human genome/transcriptome (GENCODE GRCh38) with up to 2 mismatches (31,60,61). dbSNP being such a huge database, real-time fetching of all this information about crRNAs and flanking primer sequences can take time. To reduce this lag, the extracted information regarding target clinically relevant SNPs, crRNAs, gene ID, and primer sequences, is framed as a SQLite database to support the webserver, **Figure 2**.

**Figure 2.**
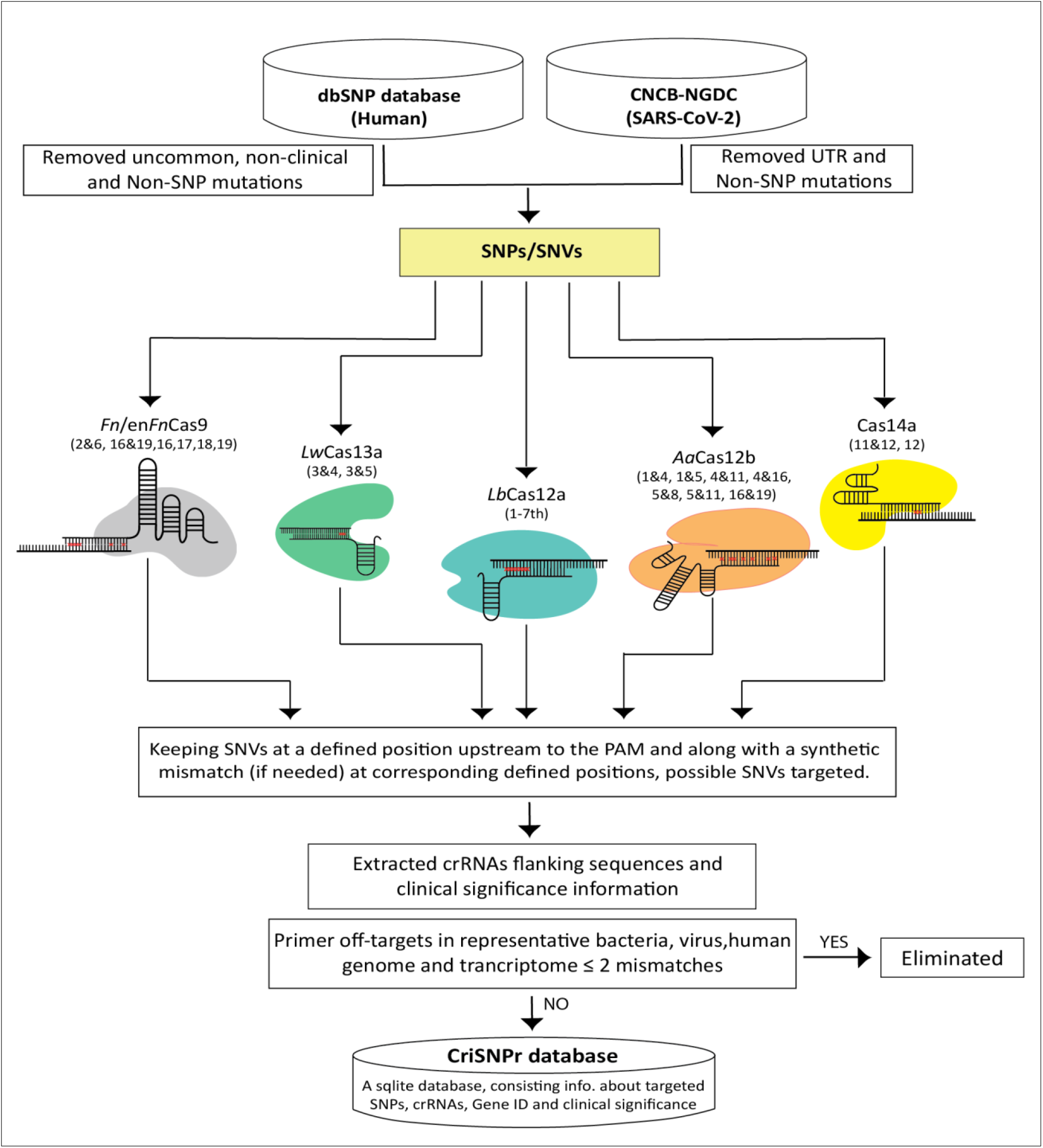
Schematic representation for CriSNPr database curation, Clinically relevant dbSNP variations were filtered for uncommon and non-SNP mutations, and non-UTR SNVs were filtered for SARS-CoV-2. The filtered SNPs were then checked for the extent of targetability by individual Cas systems based on previously reported mismatch sensitivity with or without PAM. Genome coordinates of targe SNPs assisted with acquiring information about gene IDs along with SNP flanking sequences for oligo synthesis, creating an SQLite database.

### CriSNPr interface provides a search and select pipeline for planning a CRISPRDx assay

CriSNPr’s interface has been divided into three subdomains featuring, i) Human, with ready-made target clinically relevant SNPs from human dbSNP database, ii) SARS-CoV-2, with ready-made target SNVs within SARS-CoV-2 genome, and iii) Seq-CriSNPr, to be used for sequence-based search of any reported target SNV within human or SARS-CoV-2 genomes. CriSNPr web server first tries to authenticate the entered input as a valid input by mapping and visualizing SNVs/SNPs, for related information like an organism of origin, genome coordinates, gene ID, disease association, reported allele, and their population-specific frequency distribution. The targetability of a queried SNP/SNV by all included Cas systems is simultaneously checked and only the Cas systems with positive results are shown with the required information.

The interface for human variants in the CriSNPr web server uses SNP rsID as the input and with a valid ID, looks for available matching sequences into the CriSNPr database of target SNPs, giving output as SNP related allelic information, crRNAs, and primer oligos for SNP detection by the Cas systems integrated into the CriSNPr, **Figure 3a, Methods**. The SARS-CoV-2 interface takes variant amino acid positions (for example S_N501Y, S_E484K, etc.) as the input, and after finding a match in the CriSNPr database of target SNVs, it again gives out SNV identity information along with variation frequencies, crRNAs, and primer oligos for SNV detection in the variant SARS-CoV-2 lineages by the Cas systems integrated into the CriSNPr, **Figure 3a, Methods**.

**Figure 3.**
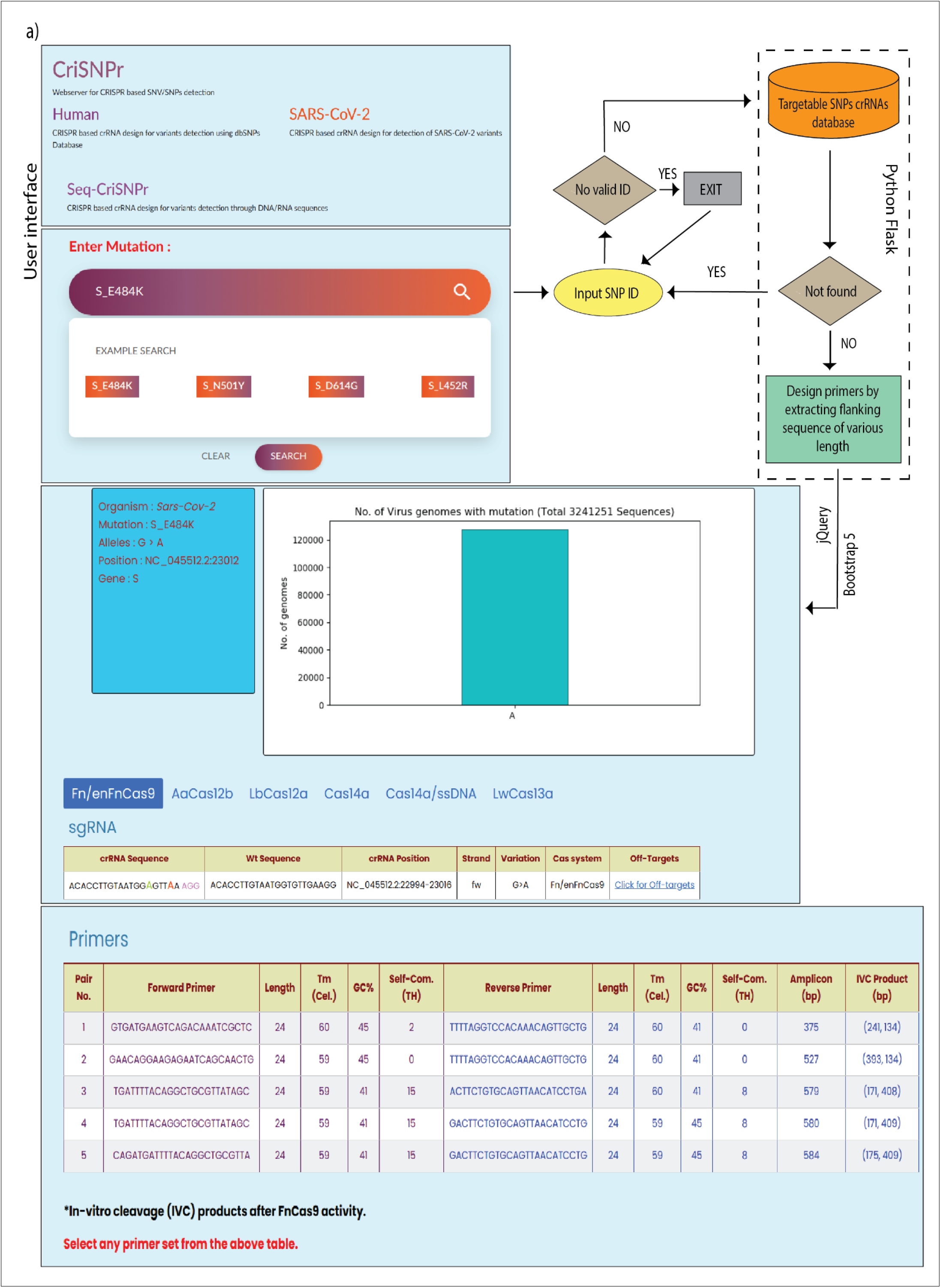
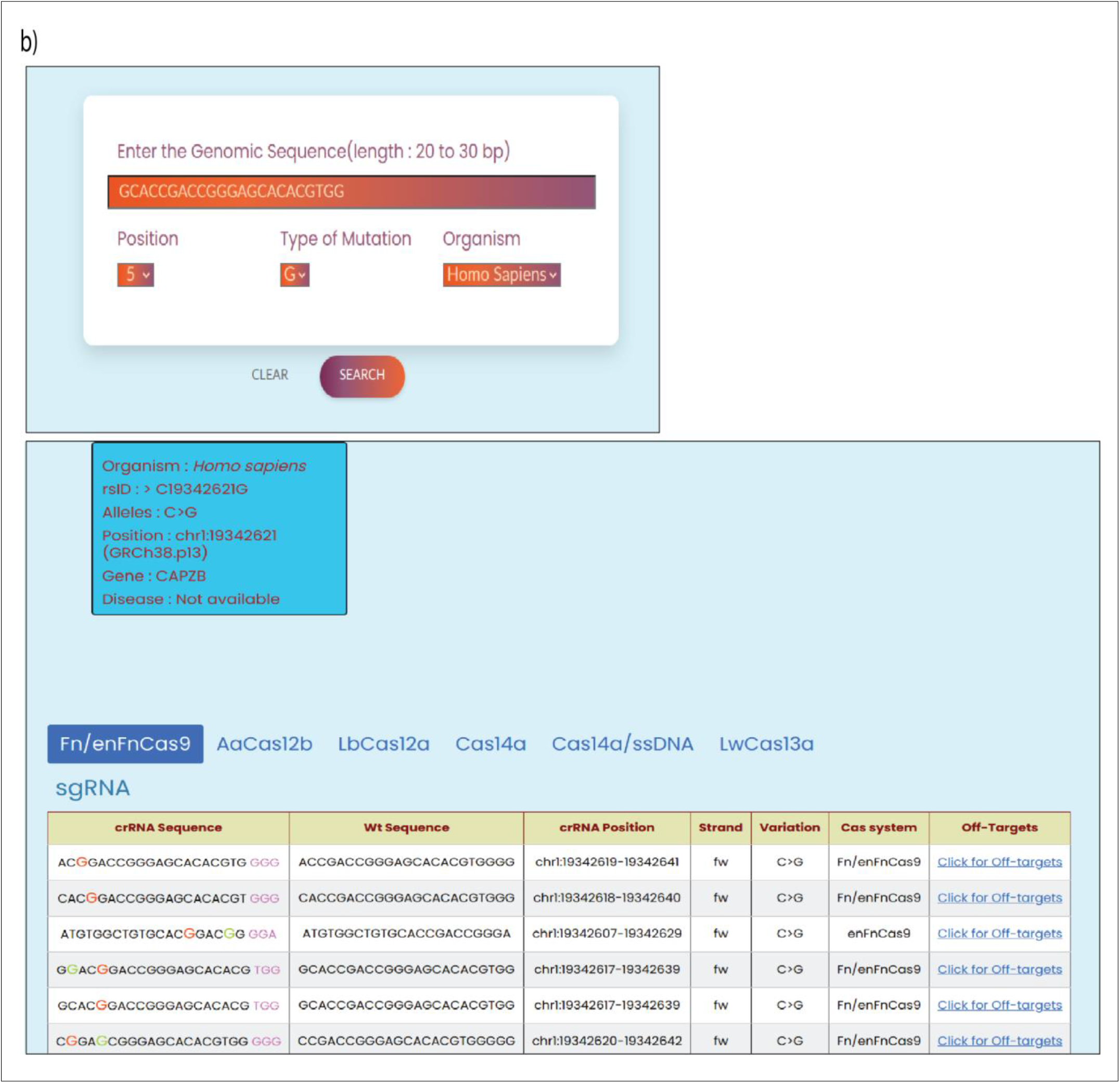
Workflow of CriSNPr webserver, **a)** CriSNPr user interface shown with three subdomains as Human, SARS-CoV-2, and Seq-CriSNPr taking rsID, mutant amino acid position, and SNV containing 20-30 nt. sequences as respective inputs. With a valid input, the server will find matching crRNA sequences in the database made by using the python flask framework. Primer sequences for amplification and SNV detection along with SNV allelic distribution, crRNA sequences, and off-targets are returned as output. **b)** Seq-CriSNPr, provides the flexibility to enter any 20-30 nt. sequences related to the human or SARS-CoV-2 genome, along with SNV nucleotide position and identity in the query sequence to give out sequence results. Representative examples are shown for user inputs.

Since the position of mismatches differs between different Cas proteins used for designing the diagnostic pipeline, CriSNPr provides also returns information highlighting the position of mutant (red color) and synthetic nucleotide (needed depending on the Cas systems used, green color) within a crRNA sequence along with the genome coordinates of the target the DNA strand corresponding to the crRNA of interest as well as the off-targets of the modified crRNA sequences. To identify potential off-targets against the modified crRNAs, CriSNPr takes the advantage of the previously reported offline versatile algorithm Cas-OFFinder (42). Using this, CriSNPr gives out off-target information against a crRNA as chromosome location and coordinates of the off-targets, DNA strand information, and a number of off-target sequences with up to 4 nucleotide mismatches, etc, **Figure 3a**. All the primer sequences provided by CriSNPr are pre-filtered for off-targets with up to 2 nucleotide mismatches against representative bacterial genome database (NCBI), virus genome database (NCBI), and human genome/transcriptome (GENCODE GRCh38). This is done particularly to prevent off-target amplification when the sample consists of a mixture of human and pathogenic nucleic acids (such as in SARS-CoV-2 infection). Considering SNV detection with clinical relevance or diagnosis, only the primer sequences with very high specificity to minimize possibilities of non-specific amplicons that can be generated from different species RNA contamination, are provided by the CriSNPr, **Figure 3a**. In addition, for Cas proteins without trans-cleavage activity (such as *Fn*Cas9/en*Fn*Cas9) *in vitro* cleavage-based discrimination is facilitated by designing primers for longer amplicons. These upon CRISPR mediated cleavage can be resolved on an agarose gel (17,20).

Since the previous two interfaces provide information curated within the CriSNPr database, the outputs can be generated in under a minute. To target any previously unreported or novel SNP or SNV present in human or SARS-CoV-2, Seq-CriSNPr considers a 20 to 30 nucleotides sequence containing the SNP/SNV along with the position and nucleobase identity of SNP/SNV, as input and does a real-time design of crRNA and primer oligos pertaining to each CRISPR/Cas detection assay, **Figure 3b, Methods**. This process can take a few extra seconds (30-40 sec.) when compared to extraction of already existing information about clinically relevant SNPs/SNVs from the CriSNPr database. Seq-CriSNPr, using the same python flask framework as of CriSNPr, gives all the sequences information similar to CriSNPr but with more user-customizable options to choose a position as well as the identity of the SNV nucleotide within a 20-30 nt query sequence, **Figure 3b**. Currently, Seq-CriSNPr takes query sequences only related to the human and SARS-CoV-2 genome but can be upgraded to other genome sequences based on the user requirements.

### CriSNPr readout can be experimentally validated in a short period of time

We next validated the CriSNPr outputs with real diagnostic assays on two substrates differing by one mismatch (mutation corresponding to E484K signature found in multiple SARS-CoV-2 variants of concern (VOCs)). To this end, we purified three of the DNA targeting effectors *Fn*Cas9, Cas14a, and *Lb*Cas12a, and performed the diagnostic assays according to previously published protocols. The crRNA sequences for SARS-CoV-2 E484K mutation detection obtained from CriSNPr, were used to discriminate between wild type and mutant sequences through difference in fluorescence intensity generated by trans-cleavage activities of Cas14a and *Lb*Cas12a, and on a paper strip (based on affinity-based discrimination) through *Fn*Cas9.

Remarkably, CriSNPr designed gRNAs were successful in discriminating between wild-type (WT) and mutant substrates based on fluorescence intensity of reporter cleavage for both Cas14a and *Lb*Cas12a, **Figure 4, Supplementary Figure 2**. Similar discrimination between wild-type and E484K containing mutant sequences was shown by *Fn*Cas9 when used with CriSNPr derived crRNA sequences for lateral flow assays visualized and quantified using a smartphone app TOPSE, **Figure 4, Supplementary Figure 2**. Taken together, the *in vitro* validation experiments with modified crRNA designs generated by CriSNPr showed the pipeline is able to design gRNAs targeting SNVs of interest in a sufficiently short period of time, **Figure 4**. Although it is conceivable that some SNVs might require more optimization than others based on the complexity of amplification and propensity of individual Cas proteins to discriminate based on design parameters, nevertheless the elimination of manual design of gRNAs with synthetic mismatches coupled with their off-target information will aid the user to focus more on refining assay components. This can be critical when rapid design of assays are required, especially during a pandemic or community outbreak of pathogenic variants of diseases.

**Figure 4.**
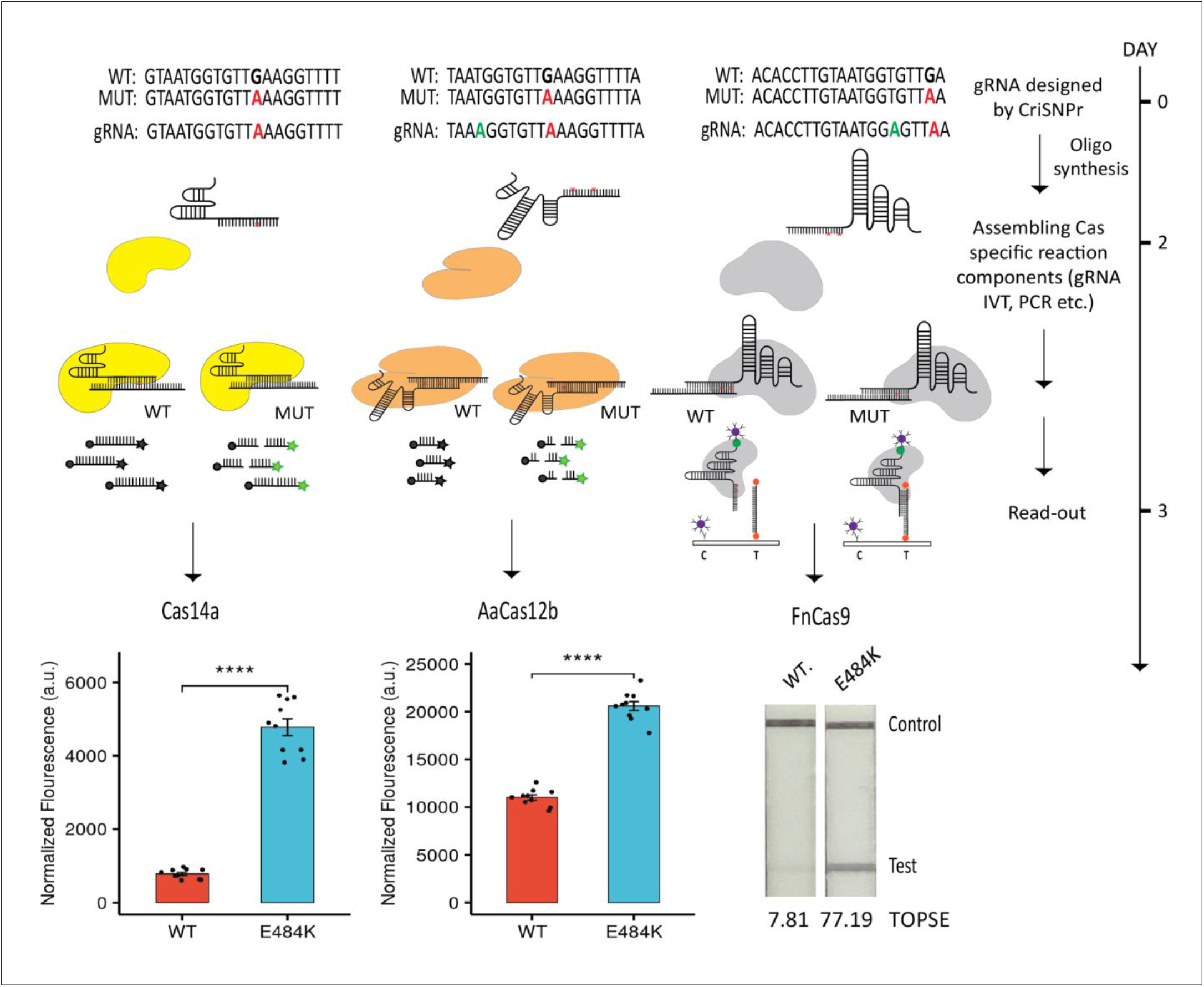
CriSNPr designed gRNAs can discriminate SNVs with multiple Cas proteins, crRNA sequences designed by CriSNPr for a representative SNV (SARS-CoV-2 E484K) detection by Cas14a, *Lb*Cas12a and FnCas9 can successfully discriminate between wild-type and mutant sequences. A tentative timeline for implementing the assays is shown on the right. Error bars represent S.E.M., student’s paired T-test p values **** ≤ 0.0001 (values from independent measurements represented as dots)

### dbSNP and SARS-CoV-2 genome targetability of different Cas systems

As described earlier, CRISPR systems showing mismatch sensitivity offered different positions for SNP targetability. To know the total target SNPs/SNVs within the dbSNP database by any of the individual Cas effectors, we further evaluated possible constraints for these systems since a systematic comparison between different platforms has not been reported so far for SNV targeting. Among the Cas proteins included, Cas13a (targeting RNA) and Cas14a (ssDNA) can naturally target almost all the variations in the dbSNP database since none of these are constrained by PAM requirements **Figure 5a**. However, Cas13a requires an additional step for converting a DNA substrate to RNA before the CRISPR reaction and readout.

**Figure 5.**
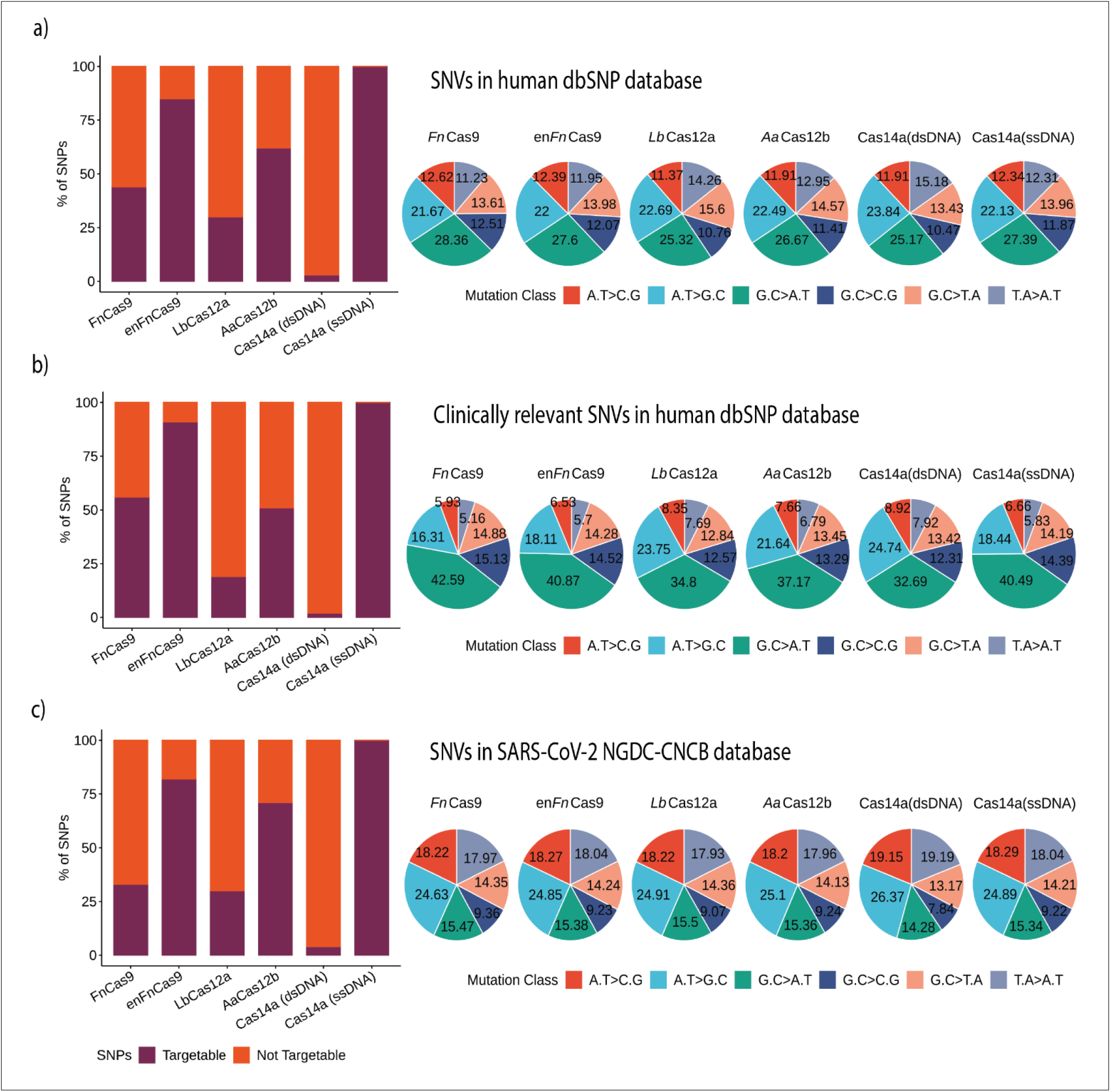
dbSNP and SARS-CoV-2 genome targetability of different Cas systems, **a)** Shows percent SNP targets for different Cas-systems across the dbSNP database along with base distribution of targeted SNPs by individual Cas-system. **b)** Percentage of targeted SNPs having clinical significance or disease relevance in humans, again with percentage base distribution at each SNP position targeted by individual Cas-system. **c)** Percentage of targeted SNPs in SARS-CoV-2 genomes reported in GISAID Database, again with percentage base distribution at each SNP position targeted by individual Cas-system. In all bar plots, red denotes non-targeting SNV percentage while violet indicates the percentage of SNVs that can be targeted.

Among the Cas systems with PAM requirements, the enhanced version of *Fn*Cas9 (en*Fn*Cas9) showed the highest numbers of target SNVs (85.15%, 15988454 SNPs). This is followed by *Aa*Cas12b, *Fn*Cas9 and *Lb*Cas12a with 62.32% (11702008), 43.86% (8234812) and 29.95% (5623279) target SNPs respectively. Cas14a could target 500786 variations (2.66%) out of the total 18775119 variations when the substrate is a double-stranded target DNA due to the TTTA PAM constraint. These findings suggest that even though the number of sensitive mismatch positions increases the possibilities of targeting an SNP from dbSNP, PAM relaxation provides more coverage for detecting SNVs. In contrast, *Lb*Cas12a despite 7 single mismatch sensitive positions targets only 5623279 SNPs because of a stringent TTTN PAM. When compared, *Aa*Cas12b provides more SNPs detection (62.32%) due to relaxed TTN PAM as well as various combinations of dual mismatch sites. Since Cas14a with dsDNA targets show sensitivity only for 2 positions coupled with very stringent TTTA PAM, the overall detection ratio is the lowest among all PAM-dependent SNPs detection Cas systems, **Figure 5a**.

Next, we checked the possible disease-causing/associated SNPs detected by different Cas systems. A total of 493105 disease-related SNPs were filtered from the SNPs present in dbSNP. Among PAM-dependent Cas systems, en*Fn*Cas9 and Cas14a (dsDNA) show the highest (90.74%) and lowest (1.50%) SNPs detection respectively, **Figure 5b**. Interestingly, besides *Fn*/en*Fn*Cas9, all the other PAM-dependent Cas systems showed a lower number of disease-related SNPs that can be targeted. This can possibly be due to the presence of NRG/NGR PAMs within the vicinity of the majority of these disease-related SNPs, **Figure 5b**.

This could be further explained by looking at the distribution of various mutation classes in target SNVs. Both total and clinically relevant target mutations in the dbSNP database are enriched for G.C>A.T mutation class in all Cas systems, **Figure 5a, 5b and Supplementary Table 3**.

In clinically relevant variations, G.C>A.T class is dominating the other mutation classes, **Figure 5b**. There is a clear difference in the abundance of mutational classes among the various Cas systems for clinically relevant SNVs. Thus G.C>A.T mutation classes can be targeted by Cas systems with G-rich PAMs such as *Fn/enFn*Cas9 (42.59% and 40.87%) compared to Cas systems with T rich PAM like *Aa*Cas12b, *Lb*Cas12a and Cas14a(dsDNA) (37.17%, 34.8% and 32.69%). Similarly *Aa*Cas12b, *Lb*Cas12a and Cas14a (dsDNA) are enriched for A.T>G.C classes compared to the *Fn/enFn*Cas9, **Figure 5b**. Since Cas14a (ssDNA) can target almost 100% SNVs available both at dbSNP and ClinVar, the mutation classes shown for Cas14a (ssDNA) can serve as reference for the other Cas systems in each case.

Considering the ongoing COVID-19 pandemic and emergence of rapidly mutating SARS-CoV-2 variants, CriSNPr also includes targets for SARS-CoV-2 SNV detection. SARS-CoV-2 variation database from CNCB-NGDC (based on GISAID genome sequences) was used as a reference to build the required database of CriSNPr. Since the SARS-CoV-2 genome is AT-rich (62.05%) compared to the human genome, which is even reflected by decreased targetability for G-rich *Fn/enFn*Cas9 and slightly higher targetability (30% and 70%) for *Lb*Cas12a and AaCas12b, respectively, **Figure 5c and Supplementary Table 4**.

## Discussion

In this manuscript, we report a single web server providing the user with pre-designed guide RNAs and flanking sequences to formulate a CRISPR-based diagnostic pipeline with ease. The field of CRISPR diagnostics has exploded in recent years with multiple Cas systems showing tremendous promise in reading and detecting nucleotide modifications in a substrate (14–20,24). As this opens up applications in multiple spheres of biotechnology and clinical diagnostic regimens-particularly for early, on-site detection, the necessity to develop design parameters that are streamlined for assay design is high. CriSNPr is one of the first web servers focused on reducing the time and effort required for designing CRISPR/Cas-based SNV detection assays. When combined with different readout modalities tailored for multiple CRISPR effectors, it can enable the design of CRISPR diagnostics for rapid detection of monogenic and infectious diseases for different Cas systems.

While this manuscript was being prepared, a few web-servers that design gRNAs specifically for targeting SNVs across the genome, especially in an allele-specific manner have been reported in the literature (62–65). These present an important advancement towards gRNA design for precision medicine in general. However, the CriSNPr platform is tailored for generating gRNAs specifically for diagnostics taking into account the design parameters for each diagnostic CRISPR protein. Thus, it caters to applications that are not covered by general gRNA design databases and toolsets, **Supplementary Table 2**.

In the current version of CriSNPr, six of the widely used CRISPR diagnostic platforms have been included due to enough literature support for their design guidelines. The field of CRISPRDx being rapidly evolving, several other Cas proteins have been recently reported for detecting pathogenic DNA or RNA and their associated variants (66,67). As more and more literature supporting these diagnostic pipelines become available, we would include them in the server. The addition of multiple CRISPR pipelines in an integrated server caters to two overarching purposes. Firstly, an SNV of interest might be detected only by specific Cas proteins and not by others, **Supplementary Figure 3**. Secondly, where multiple diagnostic options are available, users can flexibly decide the CRISPR platform of choice based on the availability of reagents and methodologies. The latter is particularly useful since the sensitivity and scope of readout modes might vary based on the diagnostic query - pathogenic polynucleotide or single nucleobase variant (monoallelic or biallelic) (68,69). While lateral flow readouts have been demonstrated for detecting full length sequences and variants, detecting carriers of disease mutations (monoallelic SNVs) is better suited to more sensitive fluorescence based readouts (14–18,20,23,70). Such decisions can be facilitated by CriSNPr recommendations for the given SNV of interest. Importantly, a wide variety of design options allows users to test and standardize the most optimum pipeline for the SNV of choice. This is particularly true since the different CRISPR systems with PAM requirements have an affinity towards AT/GC-rich PAM sequences. As the number of such sequences varies between humans and other pathogenic genomes the ability of CRISPR proteins to discriminate between SNVs vary as well based on the target species.

There are several areas where CriSNPr requires immediate improvement. For example, the current framework of CriSNPr is unable to perform batch processing of sequences. This limitation is due to the technical parameters of the system that hosts the server and will be updated in near future to accommodate batch processing. Also, CriSNPr doesn’t currently include variants that are not single nucleotide changes. Although SNVs perhaps represents the most sensitive and critical diagnostic challenge, later versions of CriSNPr will involve gRNA design parameters for polynucleotide changes in the target.

In several cases, the design strategy implemented for allocating mismatches in gRNAs will be greatly improved upon further experimental validation of gRNA mismatches by multiple Cas species. This is particularly true for several of the Cas effectors considered in this server such as *Fn*Cas9, Cas14, and Cas13 where systematic dissection of every combination of nucleobase mismatch on gRNA sensitivity has not yet been reported (15,17,18,20,23,24). As and when such data becomes available, it will be possible to enable scoring of gRNAs returned as output so users can reduce the time taken for validating more efficient pipelines.

CriSNPr currently hosts genomic targets for humans and SARS-CoV-2 lineages. This is a limitation considering that a large number of pathogenic variants exist in the microbial community and several of these have important clinical manifestations such as drug or antibiotic resistance. Although the webserver has an additional *de novo* design functionality, we hope to increase the scope of the tool to cover as many sequences that can be fetched from public repositories as possible. Perhaps the most critical evaluation of CriSNPr can be done once there are more web servers that are made available for comparison. Till that happens, CriSNPr in its current form should cater to implementing CRISPR diagnostics in a wide range of clinical and academic settings and reduce the time and effort required for design and validation for every nucleotide variant.

## Acknowledgements

We thank all members of Chakraborty and Maiti labs for helpful discussions and valuable insights pertaining to this work. This study was funded by CSIR-Sickle Cell Mission (HCP0023) and EMBO Young Investigator award, a Lady Tata Young Investigator award (GAP0198) to D.C.

## Author’s contribution

D.C., S.M., A.H.A., and M.K. conceived the project and designed the experimental pipeline. A.H.A. performed bioinformatics analysis with inputs from M.K. M.K. and S.S. performed wet-lab validation experiments. D.C., A.H.A., and M.K. drafted the manuscript with inputs from others.

## Ethics declarations

The authors declare no competing interests pertaining to this work.

## Supplementary information

**Supplementary Figure 1.**
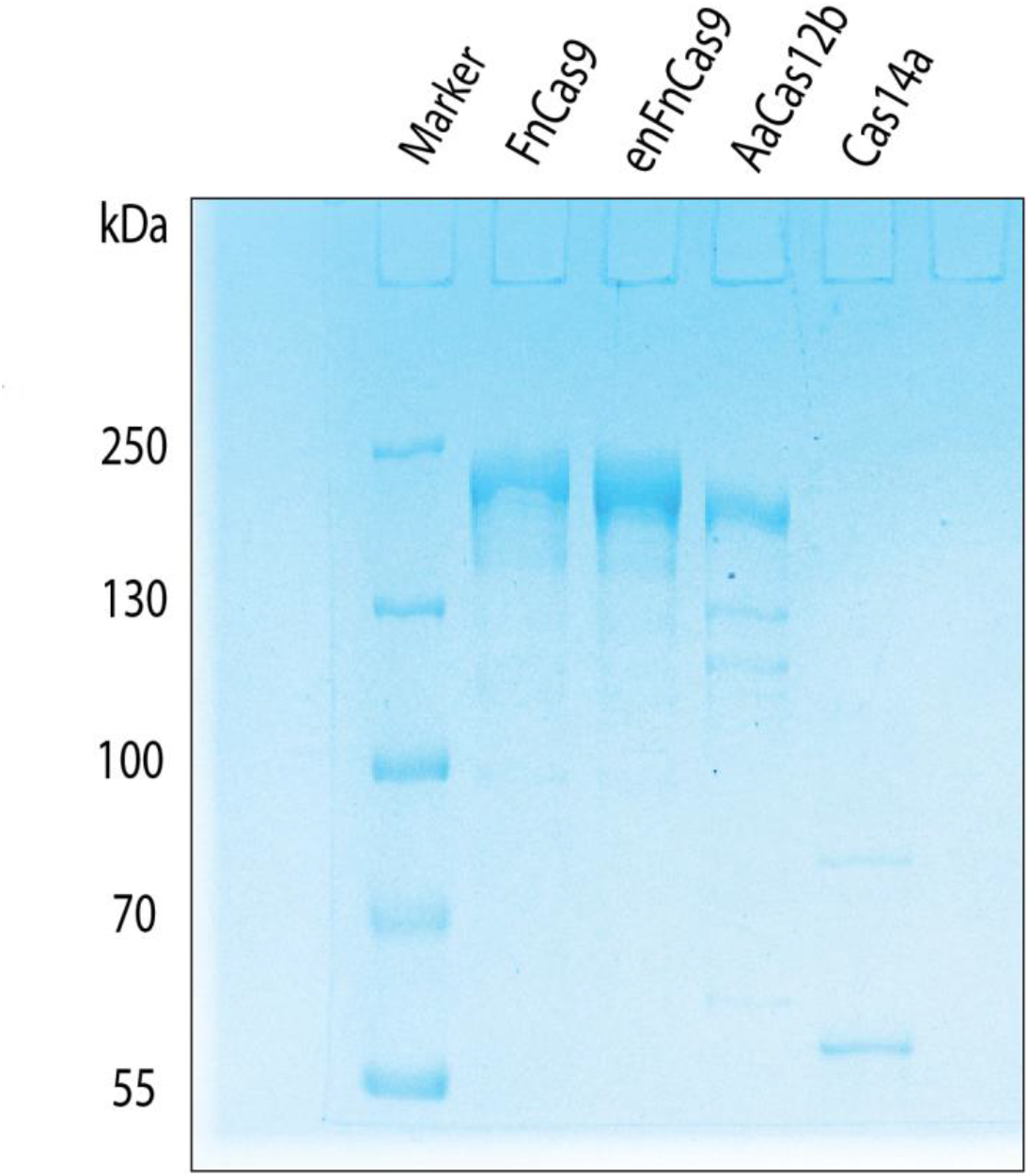
PAGE gels for purified *Fn*Cas9 (~190kDa), *enFn*Cas9 (~190kDa), *Aa*Cas12b (~130kDa) and Cas14a (~61kDa) proteins.

**Supplementary Figure 2.**
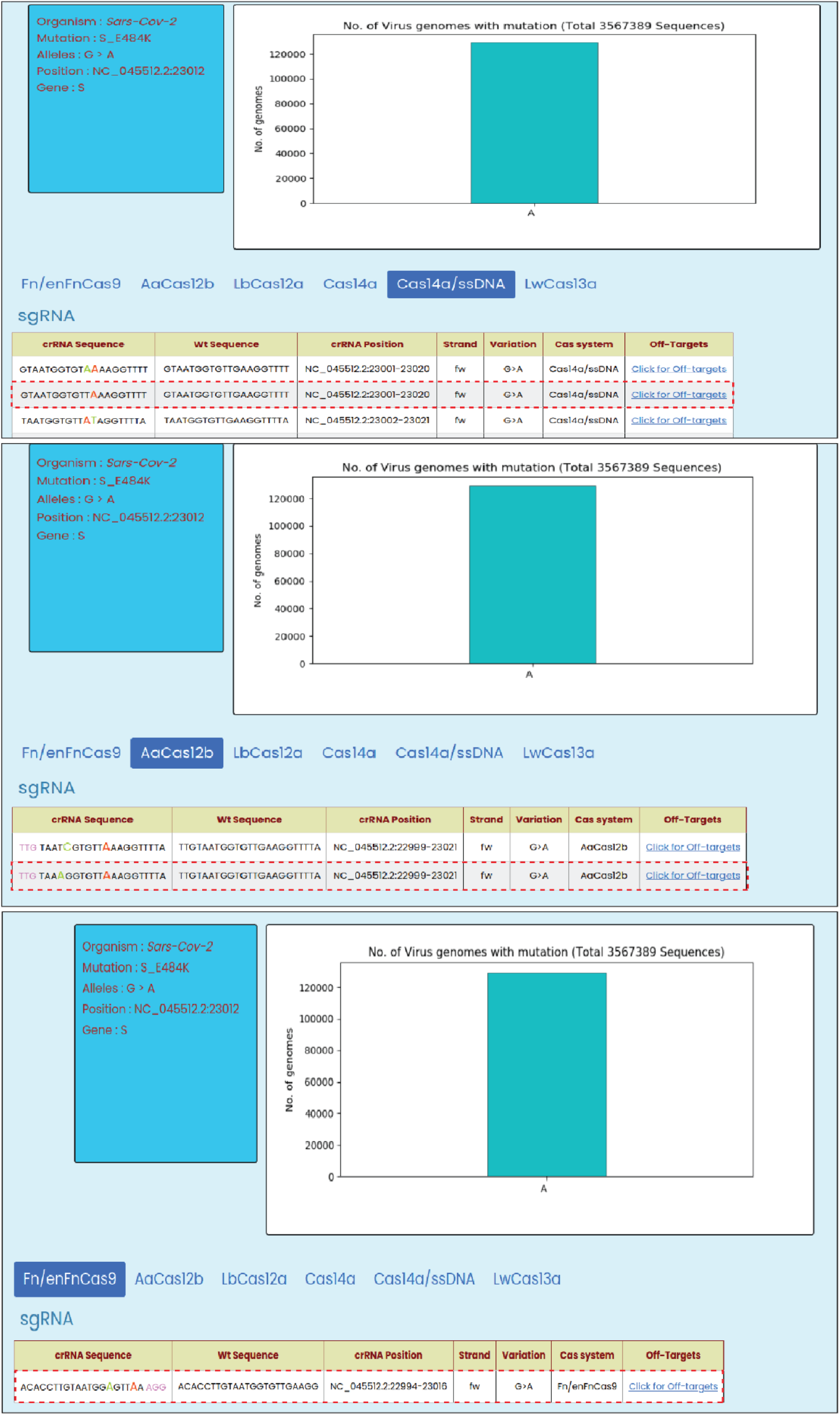
gRNA sequences designed by CriSNPr for the detection of SARS-CoV-2 E484K mutation using Cas14a, *Aa*Cas12b, and *Fn*Cas9 respectively shown in Figure 4 (denoted with red dotted box).

**Supplementary Figure 3.**
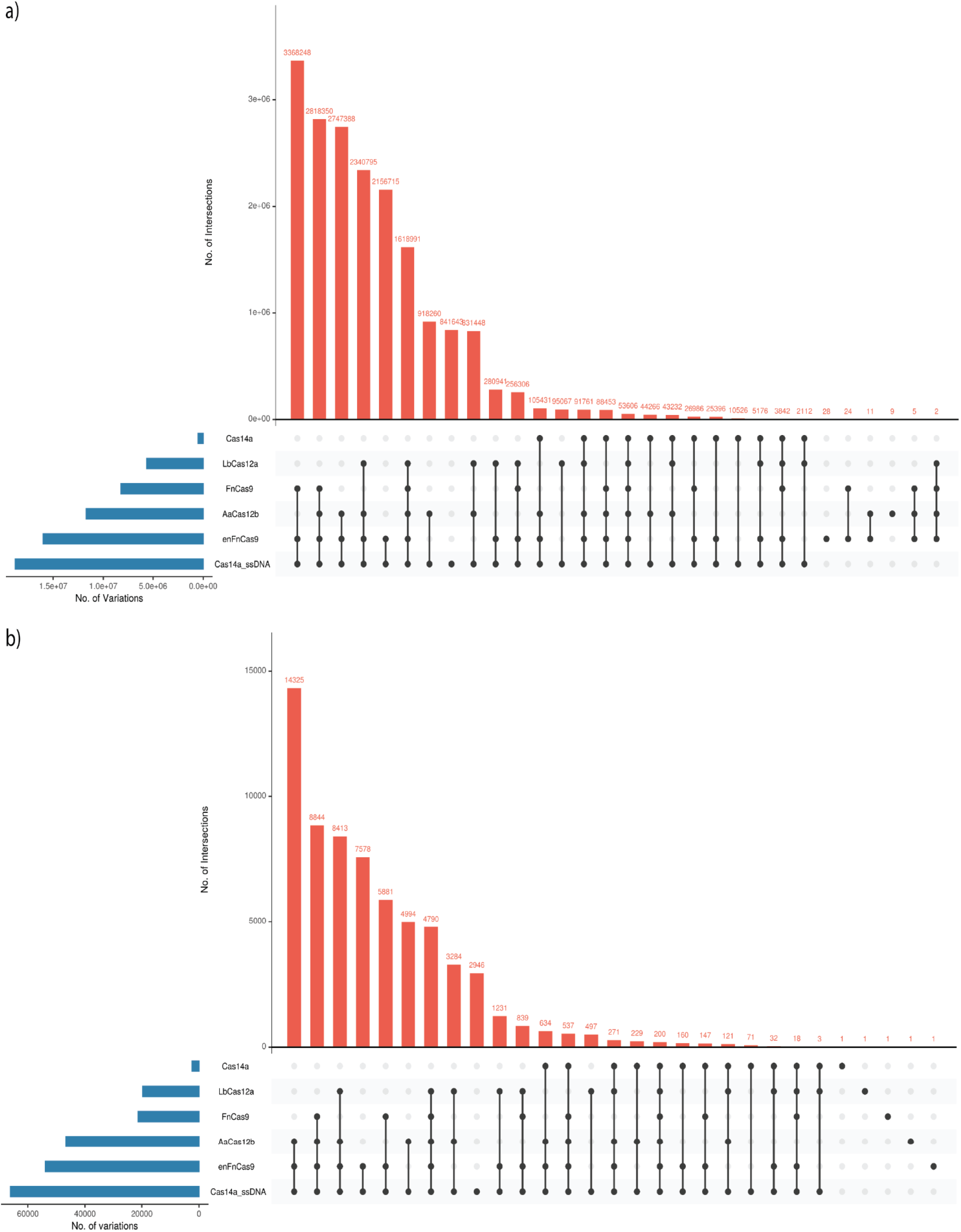
Upset plot of the intersection of the targetable variation of various Cas systems a) Human dbSNPs targetable variations b) SARS-CoV-2 CNCB-NGDC variations

**Supplementary Table 1**, List of oligos used in this study

**Supplementary Table 2**, Comparison between different sgRNA designing tools for SNVs in a sequence

**Supplementary Table 3**, Variation statistics from human dbSNP database for various Cas systems

**Supplementary Table 4**, Variation statistics from SARS-CoV-2 CNCB-NGDC database for various Cas systems

